# Genome-Wide Association Study Reveals First Locus for Anorexia Nervosa and Metabolic Correlations

**DOI:** 10.1101/088815

**Authors:** E. L. Duncan, L. M. Thornton, A. Hinney, M. J. Daly, P. F. Sullivan, E. Zeggini, G. Breen, C. M. Bulik

## Abstract

Anorexia nervosa (AN) is a serious eating disorder characterized by restriction of energy intake relative to requirements, resulting in abnormally low body weight. It has a lifetime prevalence of approximately 1%, disproportionately affects females^1,2^, and has no well replicated evidence of effective pharmacological or psychological treatments despite high morbidity and mortality^2^. Twin studies support a genetic basis for the observed aggregation of AN in families^3^, with heritability estimates of 48%-74%^4^. Although initial genome-wide association studies (GWASs) were underpowered^5,6^, evidence suggested that signals for AN would be detected with increased power^5^. We present a GWAS of 3,495 AN cases and 10,982 controls with one genome-wide significant locus (index variant rs4622308, p=4.3x10^−9^) in a region (chr12:56,372,585-56,482,185) which includes six genes. The SNP-chip heritability 
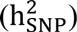
 of AN from these data is 0.20 (SE=0.02), suggesting that a substantial fraction of the twin-based heritability stems from common genetic variation. Using these GWAS results, we also find significant positive genetic correlations with schizophrenia, neuroticism, educational attainment, and HDL cholesterol, and significant negative genetic correlations with body mass, insulin, glucose, and lipid phenotypes. Our results support the reconceptualization of AN as a disorder with both psychiatric and metabolic components.

---

Following uniform quality control and imputation using the 1000 Genomes Project (phase 3)^7^ in 12 anorexia nervosa (AN) case-control cohorts, we performed association analysis using an additive model on the dosage data for each cohort and an inverse-variance weighted meta-analysis across cohorts (see **Supplementary Text** for methods, quality control details and see **Supplementary Table S1** for individual study details). Results were obtained for 10,641,224 SNPs and insertion-deletion variants with minor allele frequency > 1% and imputation quality scores > 0.6 (see Supplementary Figure S1 for quantile-quantile plot). GWAS statistic inflation (*λ*) was 1.045, and the sample size adjusted *λ*_1000_ was 1.008, suggesting minimal inflation due to population stratification or other systematic biases.

One locus achieved genome-wide significance for AN (Figure 1). Information for the top ten loci is given in **Supplementary Table S2.** The chromosome 12 (12q13.2) locus reported here is multigenic, overlaps six genes (*IKZF4*, *RPS26*, *ERBB3, PA2G4, RPL41*, and *ZC3H10*), and is located near six additional genes (*ESYT1, SUOX*, *RAB5B, CDK2*, *PMEL*, and *DGKA*). Supplementary Figure S2 provides a forest plot and information about effects across cohorts for the top SNP (rs4622308, P=4.3x10^−9^, OR (C allele) =1.2, SE=0.03, MAF_cases_=0.48, MAF_controls_=0.44), which were relatively consistent across cohorts. Results of conditional analyses are consistent with the existence of one signal at this locus (see Supplementary Figure S3). Several other immune-related phenotypes: vitiligo, alopecia areata, and asthma (see Supplementary Figure S4) have associations in the region, although these are (somewhat) LD independent. The second (rs200312312 on chromosome 5, p=6.7x10^−8^) and fourth (rs11174202 on chromosome 12, p=3.1x10^−7^) most significant loci in our analyses also have consistent evidence for association across multiple cohorts and patterns of correlated variants with similar p-values (see Supplementary Figure S5 for area plots of these loci). The fourth best locus is intronic in the *FAM19A2* gene.

**Figure 1.**
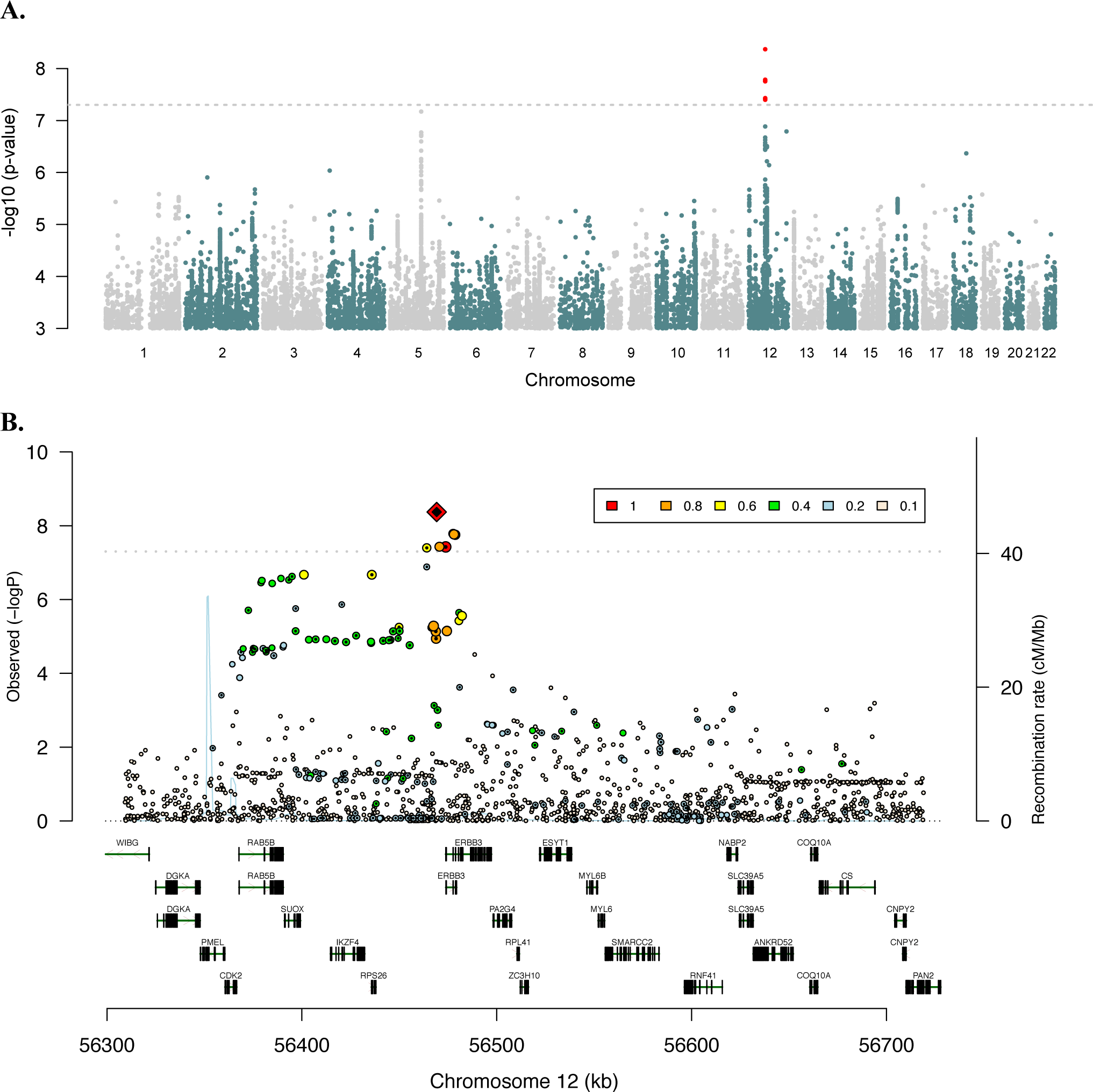
Manhattan and regional plots of the genome-wide significant locus associated with AN. A. Manhattan plot depicts a genome-wide significant locus on chromosome 12. B. Regional plot of the top locus reveals numerous genes in the region. Results depicted here reflect the full meta-analysis. Per text, see Supplementary Figure 1 for area plot with phenotypic associations. The right axis gives recombination rate, depicted with a light blue line.

Gene-wide and pathway analyses were also carried out. Multiple genes, all of which were in the region around the top SNP (rs4622308) reached gene-wide significance (reflecting the high LD in the region). Pathway analyses did not reveal experiment-wise significance (see **Supplementary Table S3** for the complete gene-wide and pathway analysis results); the top ranked pathway was CP:KEGG_LONG_TERM_POTENTIATION. As has typically been reported for other psychiatric disorders, candidate genes from previous studies did not reach significance (for a detailed review of the candidate gene literature see^8^).

Interrogation of databases such as GTeX^9^ does not indicate that any of the genes in the top region have distinct patterns of brain gene expression and searches using both GTeX and the SNP tag lookup function in MRbase (www.mrbase.org/beta) indicate that that the top SNP (rs4622308) is not, directly or via LD tagging, an eQTL or mQTL. Analyses of gene expression of the genes surrounding rs4622308 in hypothalamus from fasted and refed mice did not implicate specific genes (See Supplementary Figure S6). However, we note that rs4622308 is in high LD (r^2^=0.86; D’>0.99) with rs11171739, which has been found to be associated in GWASs of autoimmune phenotypes including type I diabetes^10^ and rheumatoid arthritis^11^. The risk associated alleles of both SNPs (C-C) are typically found on the same haplotype.

LD score regression (LDSC)^12^ was used to calculate genome-wide common variant heritability 
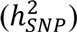
, partitioned heritability, and genetic correlations (*r_g_*) between AN and other psychiatric, medical, and educational phenotypes. In our cohort, 
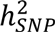
 for AN was 0.20 (SE=0.021, intercept constrained to one), comparable to 
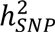
 estimates for other psychiatric disorders (Figure 2). Partitioned heritability estimates for annotation categories and cell types were not significant after multiple testing correction (for complete results see **Supplementary Table S4**).

**Figure 2.**
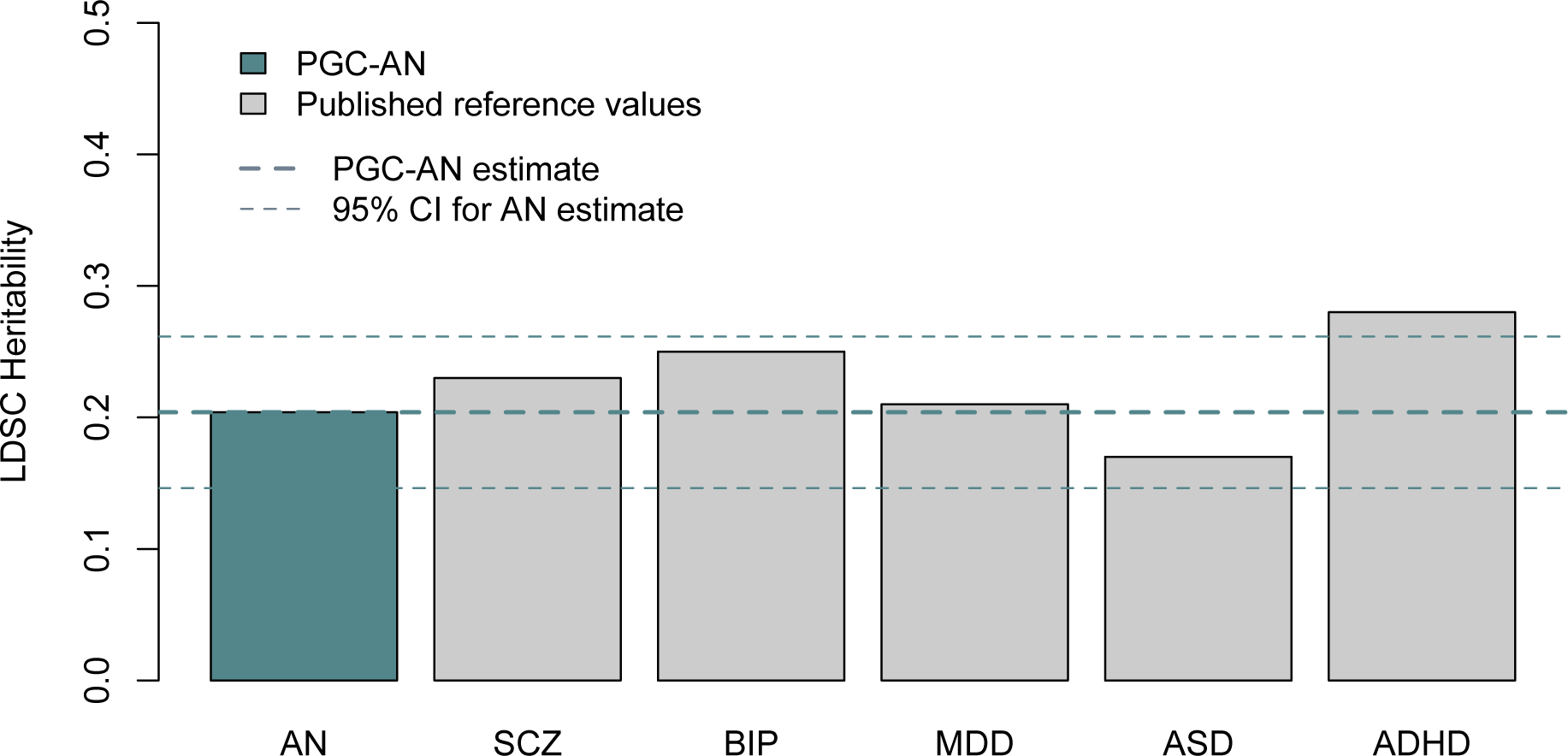
Genome-wide common variant SNP heritability estimate 
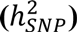
 for AN is comparable to that of other psychiatric disorders. LDSC=Linkage disequilibrium score regression, PGC-AN=Psychiatric Genomics Consortium-Anorexia Nervosa group, CI=confidence interval, SCZ=schizophrenia, BIP=bipolar disorder, MDD=major depressive disorder, ASD=autism spectrum disorder, ADHD=attention deficit hyperactivity disorder. Error bars show ±1.96xSE.

A wide range of genetic correlations between AN and other phenotypes were statistically significant. Of 159 phenotypes tested, 29 had FDR<0.05 (uncorrected p-values reported below). See Figure 3 for depiction of these genetic correlations and text below for selected examples. All 159 genetic correlations and relevant references are available in **Supplementary Table S5**.

**Figure 3.**
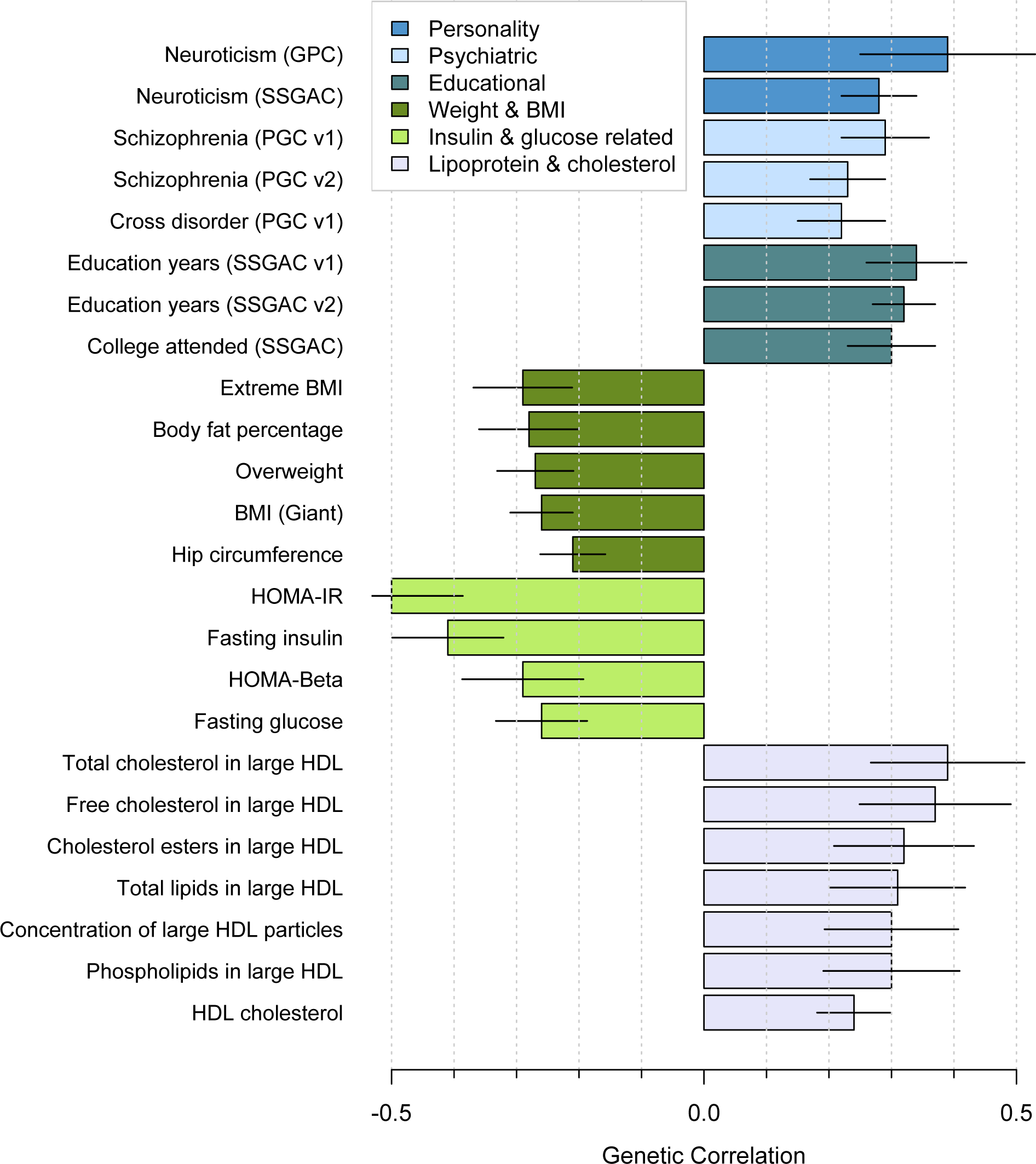
Genetic correlations between AN and diverse phenotypes reveal overlap across psychiatric, educational, weight, insulin, lipoprotein, and cholesterol phenotypes. The 24 correlations depicted here (of 159 phenotypes tested) have FDR<0.05. Bars are ± standard error.

Notable significant genetic correlations between AN and psychiatric traits and disorders included neuroticism (*r_g_*=0.39, SE=0.14, p=4.4x10^−3^), schizophrenia (*r_g_*=0.29, SE=0.07, p=4.4x10^−5^), and results from a meta-analysis across psychiatric phenotypes (*r_g_*=0.22, SE=0.07, p=3.4x10^−3^). Genetic correlations between AN and educational phenotypes such as years of education (*r_g_*=0.34, SE=0.08, p=5.2x10^−6^) and attending college (*r_g_*=0.30, SE=0.07, p=4.4x10^−5^) were also significant. Obsessive compulsive disorder (OCD) GWAS data were unavailable to us but a previous analysis has reported a *r_g_* of 0.53 (SE=0.11, SE=0.13, p=5.5x10^−6^)^13^.

Several significant negative genetic correlations emerged between AN and weight-related phenotypes, suggesting shared genetic loci underlying these phenotypes and opposing effects for relevant alleles. Extreme high BMI was significantly negatively correlated with AN (*r_g_*=−0.29, SE=0.08, p=2.0x10^−4^) as were obesity, BMI in the normal range, overweight, and hip circumference, with genetic correlations ranging from −0.2 to −0.3.

We also observed significant negative genetic correlations between AN and insulin and glucose related traits—importantly GWAS of these traits were corrected for the effects of weight/BMI: e.g., HOMA-IR (*r_g_*=−0.50, SE=0.11, p=1.3x10^−5^); fasting insulin (*r_g_*=−0.41, SE=0.09, p=5.2x10^−6^); and fasting glucose (*r_g_*=−0.26, SE=0.07, p=3.0x10_−4_). Regarding cholesterol and lipid measures, a sharp distinction between different lipid fractions is evident, for some but not all HDL vs. LDL and VLDL phenotypes. Genetic correlations between AN and HDL phenotypes were positive: e.g., total cholesterol in large HDL particles (*r_g_*=0.39, SE=0.12, p=1.6x10^−3^); free cholesterol in large HDL particles (*r_g_*=0.37, SE=0.12, p=2.2x10^−3^); and phospholipids in large HDL particles (*r_g_*=0.30, SE=0.11, p=6.7x10^−3^). In contrast, VLDL cholesterol phenotypes were negatively correlated with AN, albeit with nominal significance: e.g., total lipids in VLDL (*r_g_*=−0.30, SE=0.12, p=0.01); phospholipids in VLDL (*r_g_*=−0.33, SE=0.13, p=4.4x10^−3^); and LDL cholesterol (*r_g_*=−0.20, SE=0.08, p=0.011).

To our knowledge, this is the first robust report of a genome-wide significant association for AN. As is typical of many GWAS loci for complex disorders, the region implicated is relatively broad, and the genetic effect is relatively subtle^14^. Our genome-wide 
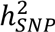
 estimate of 20% for AN supports a substantial role for common genetic variation. As we now expect^15^, the 
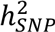
 estimate reported here indicates that common variants account for a substantial portion of twin-based heritability 
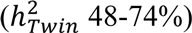
^6^. In general, 
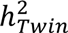
 should exceed 
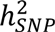
 because it captures the effects of all types of genetic variation (common and rare, as well as variation not captured with current methods).

Although genetic correlations must be interpreted in the context of multiple caveats, the observed pattern of genetic correlations with other phenotypes provides grounds for broadening our conceptualization of the disorder. First, the strong positive genetic correlations of AN with OCD and neuroticism reinforce clinical observations that high perfectionism and neuroticism predicts subsequent onset of AN^1^, which also accords with evidence of familial co-aggregation with multiple anxiety phenotypes, even when controlling for proband comorbidity^16^. Second, the positive genetic correlations seen with schizophrenia and the cross psychiatric disorder phenotype firmly anchor AN with other adult psychiatric disorders and reflect the substantial evidence for partially shared genetic risk across many psychiatric disorders^17^. Third, positive genetic correlations between AN and educational attainment suggest that genetic factors may partially account for reported associations of AN with higher familial educational attainment^18^. A positive genetic correlation between an earlier AN GWAS and educational attainment and weaker (although nominally significant) positive correlations between AN and old age general cognitive function were also previously reported^19^. Intriguingly, the association between AN and educational attainment has typically been attributed to socioeconomic factors or demands to succeed^18^; however, our data suggest potential confounding as overlapping genetic factors may mediate both.

Fourth, the identification of significant negative correlations between AN and BMI-related and anthropometric measures could potentially serve as an important first step toward gaining a better understanding of the shared biology underlying extremes of weight dysregulation (i.e., obesity vs. AN). As noted by Bulik-Sullivan et al.^12^ and Hinney et al.^20^, these results extend our understanding that the same genetic factors that influence normal variation in BMI, body shape, and body composition may also influence extreme dysregulation of these weight-related features in AN. This pattern of observations complements prior strong evidence for the involvement of neural mechanisms in obesity^21^. Finally, positive correlations with (BMI corrected) “favorable” metabolic phenotypes (i.e., HDL and lipid measures) and negative correlations with “unfavorable” metabolic phenotypes (i.e., fasting insulin, fasting glucose, HOMA-IR) encourage additional exploration of the role metabolic factors may play in extreme dysregulation of appetite and weight in AN.

Adequate explanations for how individuals with AN reach and sustain exceedingly low BMIs have been elusive. Whether the observed metabolic correlations are driven by low BMI is unknown (although most of the effects are independent of general BMI, as those GWASs corrected the phenotypes for BMI); however, future identification of specific shared risk loci and emerging analytic techniques may enable us to determine causal pathways. A physiological consequence of AN is loss of fat tissue, and similarly, there have been reports of severe metabolic changes in both acquired or congenital lipodystrophies^22^. It is possible that our final AN samples are enriched for a subgroup of patients with incipient AN who have a genetic predisposition of metabolic dysfunction, thus generating the observed genetic correlations. Additionally, the pathophysiology of AN may require two different considerations: one where the primary dysfunction is in metabolism and the eating disorder part is secondary, and vice versa.

Finally, also worthy of further investigation are the numerous immune-related phenotypic associations nearby our top locus for AN. The shared effects between AN and immune phenotypes fit into a broader pattern of above-chance comorbidity across psychiatric and immune phenotypes^23,24^. Evidence suggests that shared risk is at least partly genetic in origin^12,25^. A negative genetic correlation between AN and rheumatoid arthritis was previously reported^12^, and our LDSC estimate—though only nominally significant—is in the negative direction as well (see **Supplementary Table S5**).

The locus we identified to be associated with AN is broad and multigenic (chr12:56,372,585-56,482,185). Mechanistic explanations about the role of this variant require functional data; nevertheless, we note the possible role for genes at this locus in the pathophysiology of AN. *PA2G4* is involved in growth regulation and acts as a corepressor of the androgen receptor^26^. *ESYT1* (extended synaptotagmin-1 which binds and transports lipids^27^) is enriched in the postsynaptic density of individuals with schizophrenia^28^ and involved in calcium signaling^29^. Various studies have implicated post-synaptic density in schizophrenia pathology^30^, and calcium signaling is now believed to be etiologically relevant to schizophrenia, bipolar disorder, and autism. We speculate that the positive genetic correlation observed between AN and schizophrenia could potentially reflect these processes. Finally, rs4622308 is in high LD with autoimmune GWAS hits for type 1 diabetes^10^, and rheumatoid arthritis^11^, and the region around it harbors multiple other autoimmune associations.

In summary, we identify the first robust genome-wide significant locus for AN, which is also a previously reported type 1 diabetes and general autoimmune disorder locus. Perhaps of greater importance, is that we find AN is a complex heritable phenotype with intriguingly large and significant genetic correlations not only with psychiatric disorders but also multiple metabolic traits. This encourages a reconceptualization of this frequently lethal disorder as both psychiatric and metabolic. Just as obesity is increasingly considered to be both a metabolic/endocrine and psychiatric disorder, approaching AN as both a psychiatric and metabolic condition could ignite interest in developing or repositioning pharmacologic agents for its treatment where currently none exist.

## URLs

SNP results, https://www.med.unc.edu/pgc.

## Acknowledgments

We are grateful to the participants in these studies, without whom this work would not have been possible. CMB acknowledges funding from the Swedish Research Council (VR Dnr: 538-2013-8864). We thank all study coordinators, volunteers, and research staff that enabled this work and the Price Family Collaborative Group (PFCG). GB acknowledges support from the National Institute for Health Research (NIHR) Biomedical Research Centre at South London and Maudsley NHS Foundation Trust and King’s College London. The views expressed are those of the authors and not necessarily those of the NHS, the NIHR, or the Department of Health.

Detailed acknowledgements, funding and conflict of interest are provided in

## Author contributions

See **Supplementary Table S6**

## Supplementary Figures

**Supplementary Figure S1.**
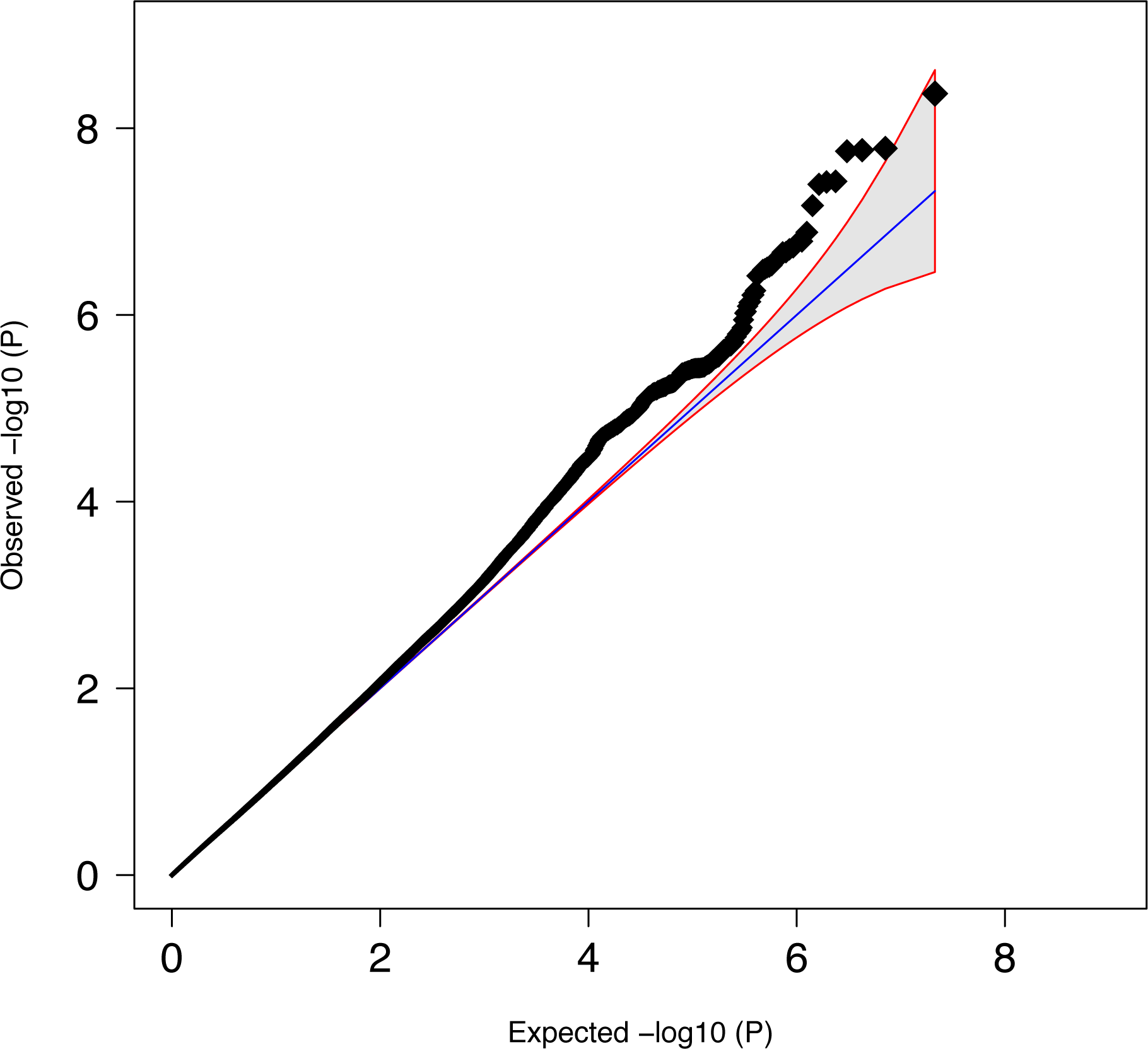
QQ plot of 10,641,224 variants with MAF>1% and imputation quality (INFO)>0.6. 15,043,779 total variants, lambda=1.045, lambda_1000_=1.008.

**Supplementary Figure S2.**
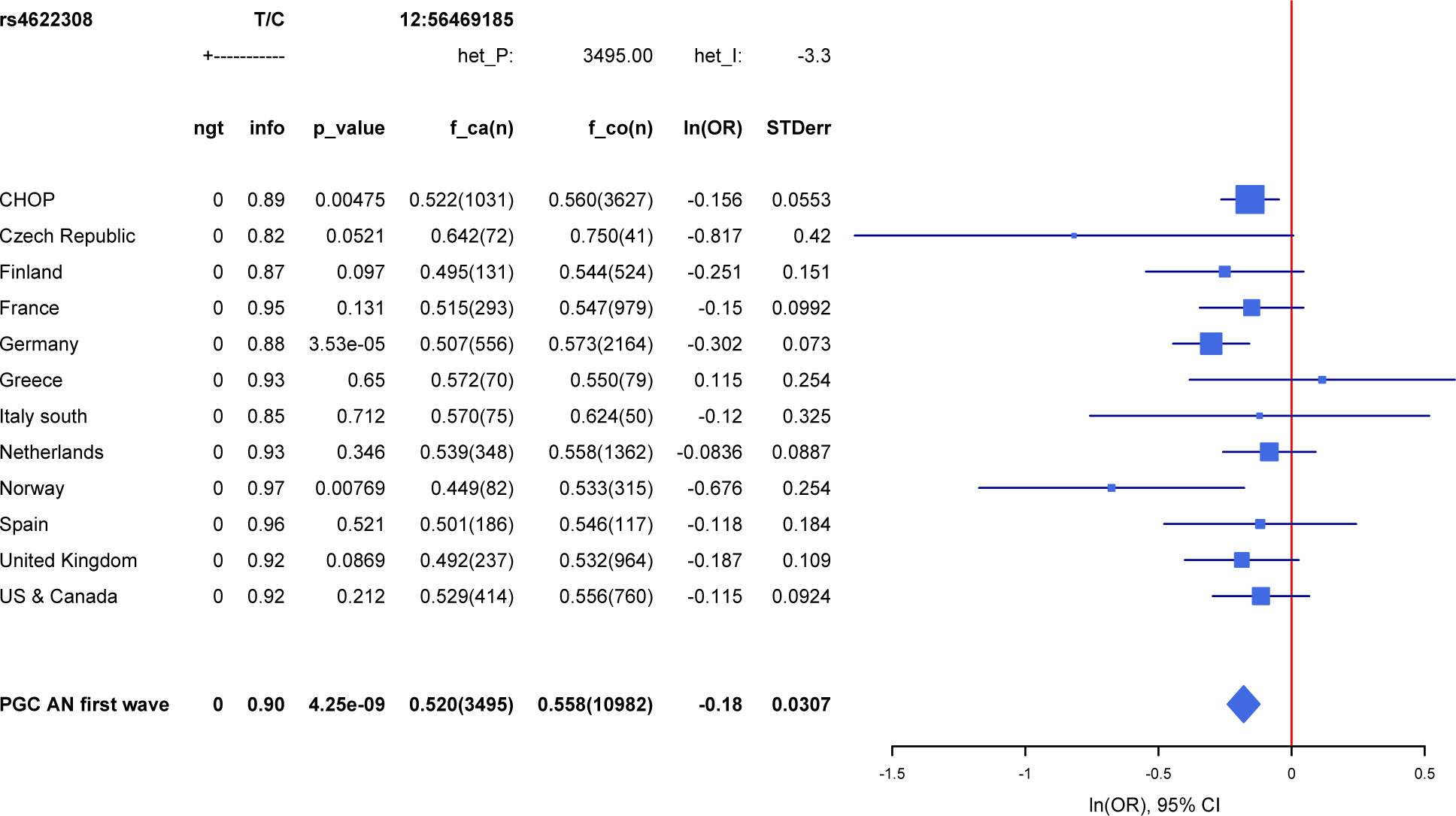
Forest plot for top SNP rs4622308. Abbreviations: ngt=number genotyped, info=imputation quality score, f_ca(n)=frequency for allele 1 in cases with number of cases in parentheses, f_co(n)=frequency for allele 1 in controls with number of controls in parentheses, ln(OR)=natural logarithm of odds ratio, STDerr=standard error.

**Supplementary Figure S3.**
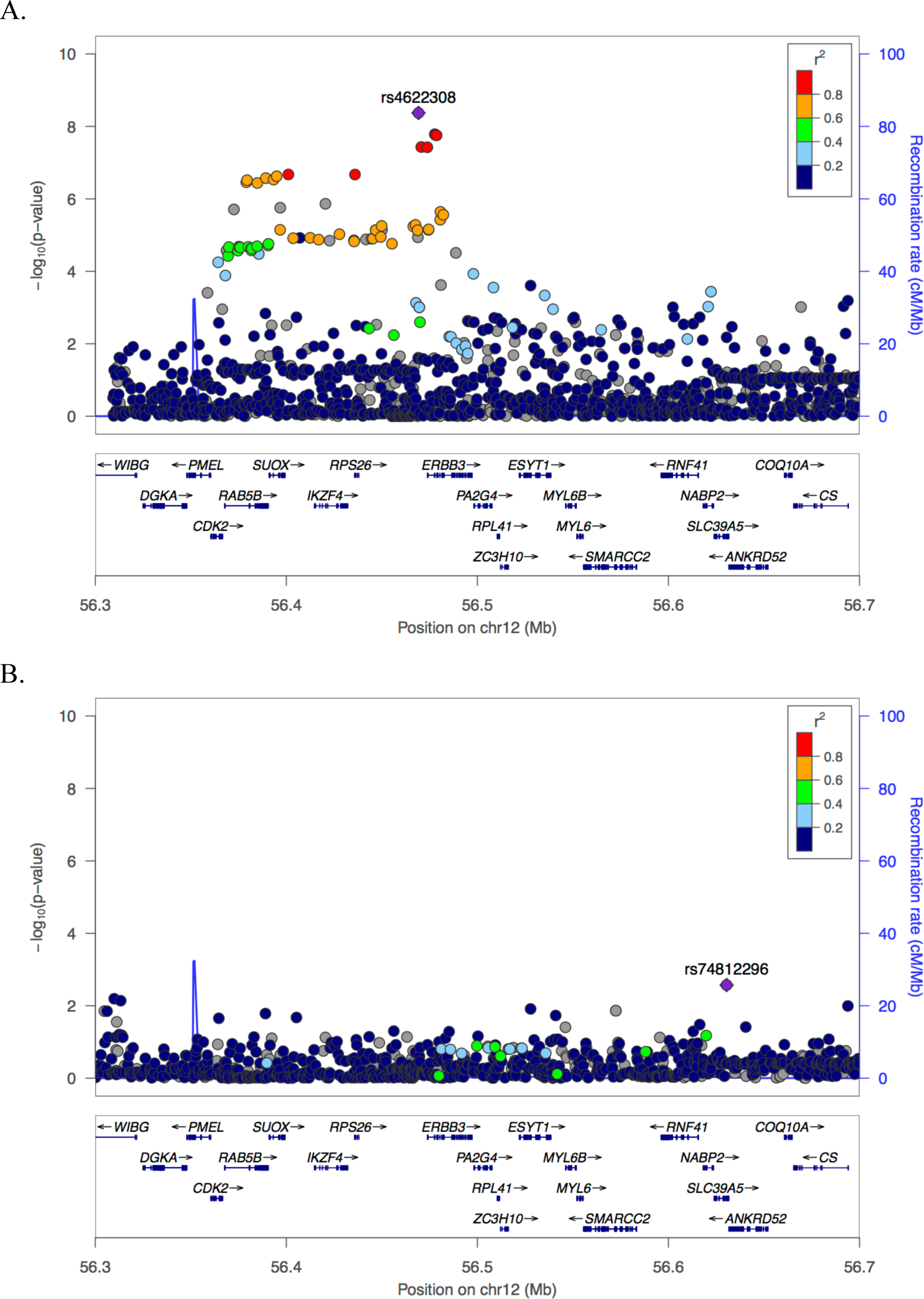
Results for top locus before and after conditioning on individuals’ dosage data at top SNP rs4622308 are consistent with the existence of only one signal at this locus. A. Before conditioning (i.e. original analysis). B. After conditioning.

**Supplementary Figure S4.**
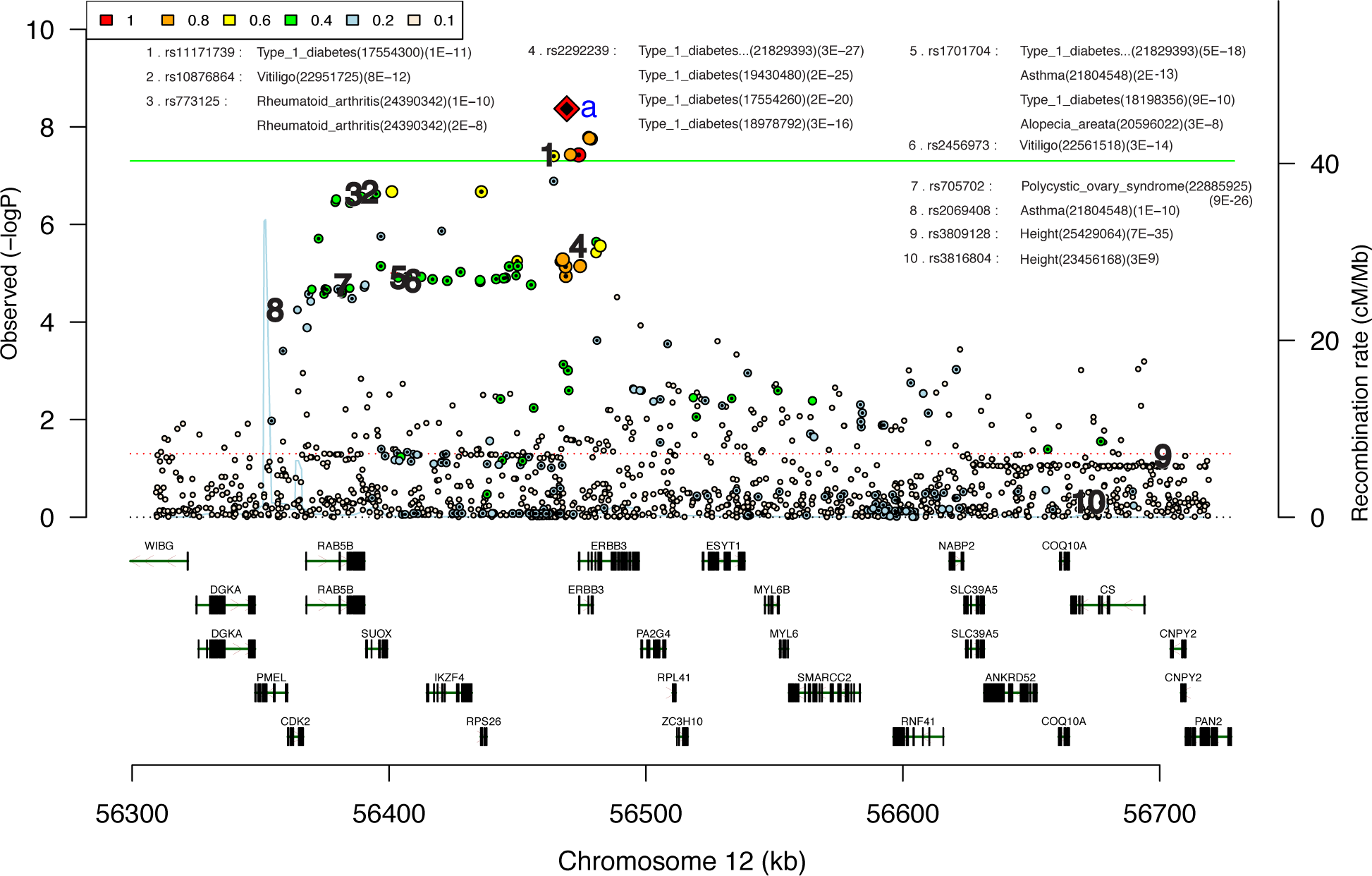
Area plot for top locus with other phenotypic associations in the region. The top SNP rs4622308 is located near phenotypic associations to numerous immune-related traits.

**Supplementary Figure S5.**
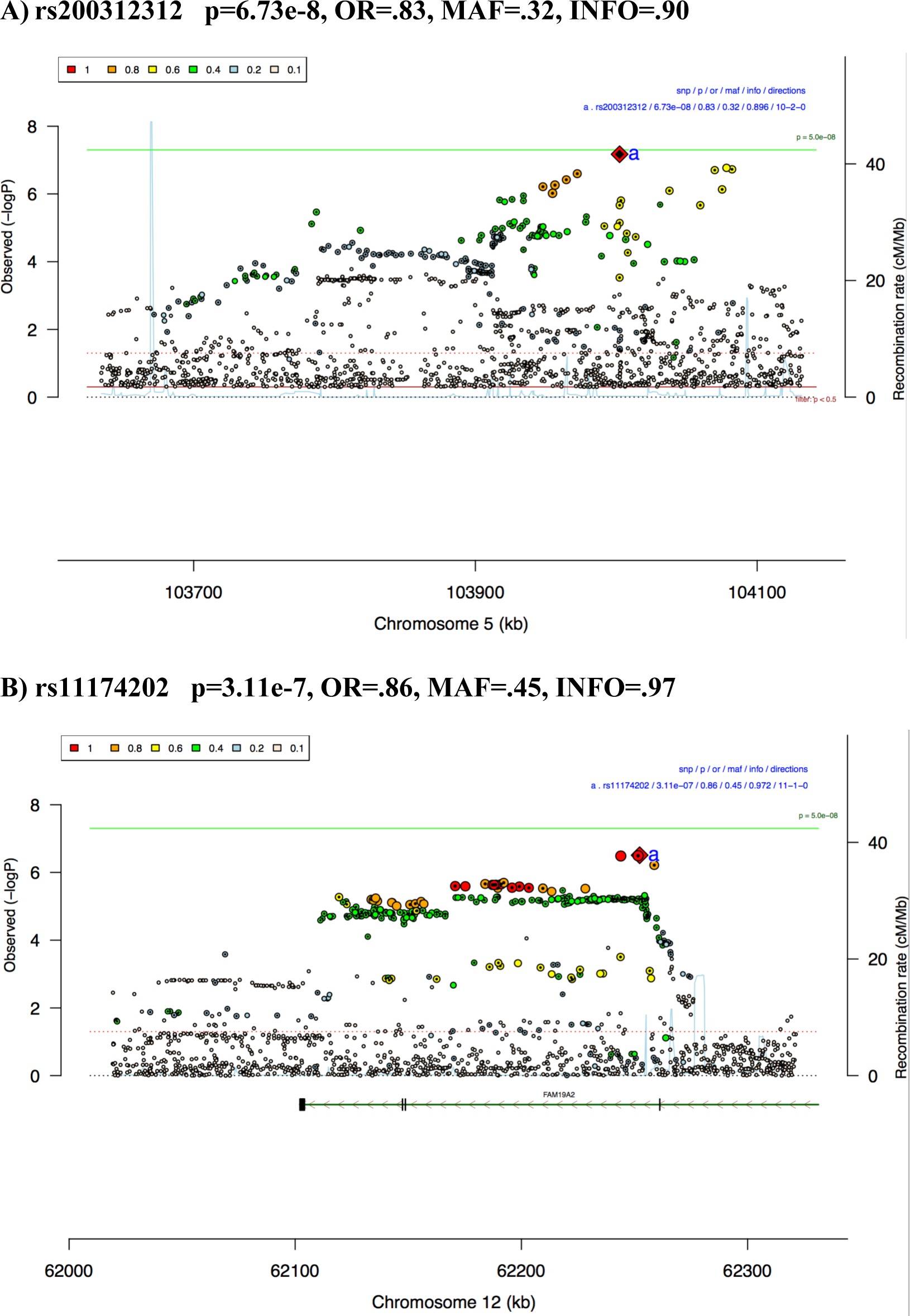
Area plots for second and fourth best loci. A) Second best locus with top SNP rs200312312. B) Fourth best locus with top SNP rs11174202. The third best locus (not shown) is a lone, rare SNP (minor allele frequency=.02), with moderate imputation quality (INFO=.7), and was consequently deemed less interesting than the 2^nd^ and 4^th^ best loci.

**Supplementary Figure S6.**
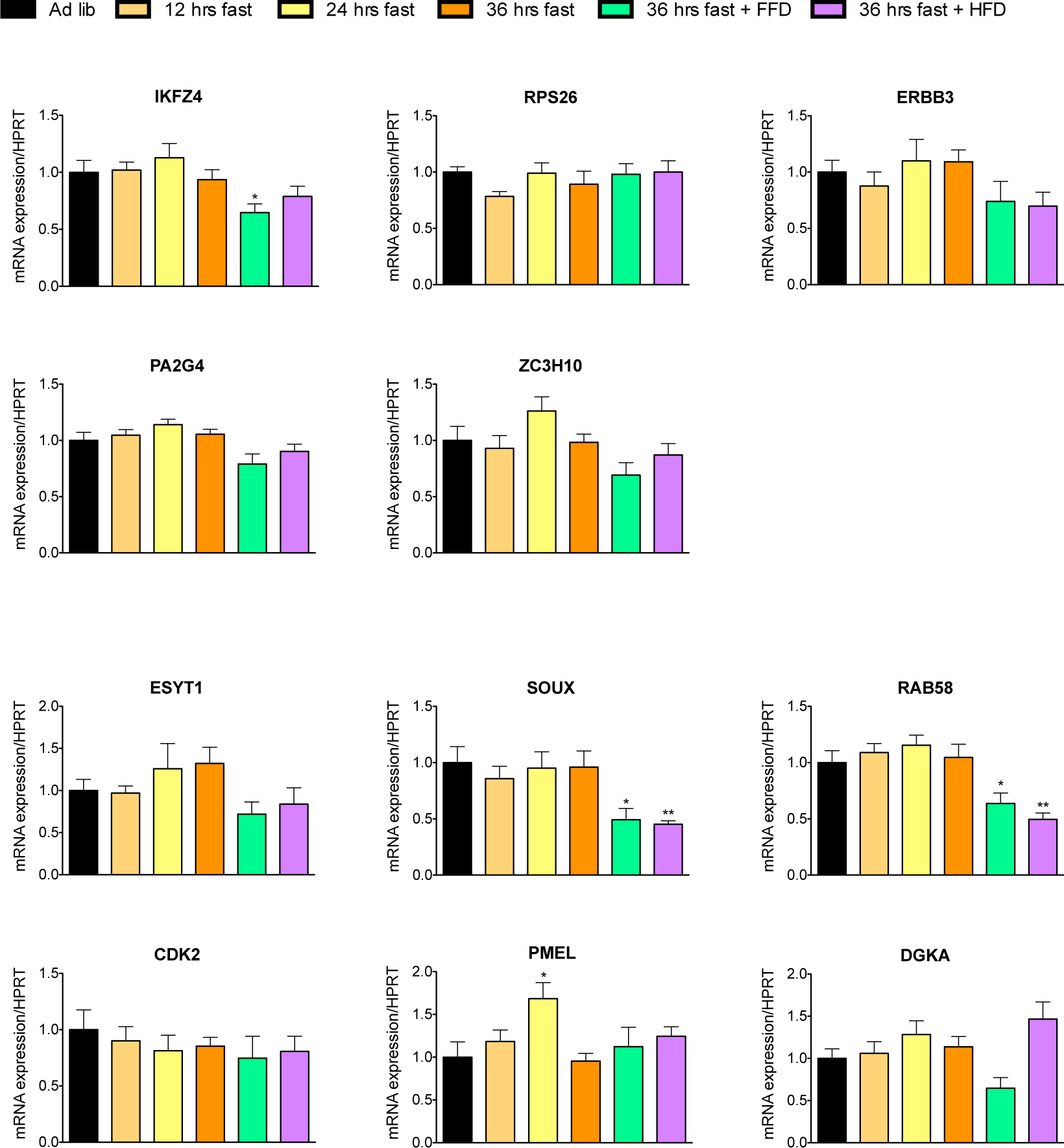
Gene expression of the genes in the rs4622308 region in mouse hypothalamus from fasted and refed C57BL/6J mice (N=5-7 for each gene). Normalised gene expression with standard errors are shown. No changes reached significance.

